# Discover Stable Expression Hot Spot in Genome of Chinese Hamster Ovary Cells Using Lentivirus based Random Integration Method

**DOI:** 10.1101/467936

**Authors:** Songtao Zhou, Xuefeng Ding, Lei Yang, Yun Chen, Xiaohai Gong, Jian Jin, Huazhong Li

## Abstract

Traditional construction method for stable expression cell line was mainly achieved by using random integration method. However, target gene might be integrated into heterochromatin region or unstable region of chromatin by using this method, and thus required multiple rounds of selection to obtain desirable expression cell lines with stable expression level of target proteins. Rational cell line construction method can overcome this shortcoming by integrating transgenes into stable hot spot within genome specifically. Thus, to discover novel effective hot spot would be critical for this new cell line construction method. Here we reported one practical method to discover new stable hot spot by using lentivirus infection and random integration. One such hot spot located at NW_006880285.1 was thoroughly investigated in this article. The expression stability of this hot spot was verified by detecting Zsgreen1 reporter gene expression for over 50 passages. When cells were adapted to suspension culture, they successfully kept expressing Zsgreen1 reporter gene. In addition, this suspension cell line could keep expressing reporter gene stably for another 50 passages. In summary, this research offered an easy new method for researchers to identify their own stable hot spots within CHO genome, and would contribute to the development of site-specific integration study in the future.

## 1. Introduction

For over three decades, Chinese Hamster Ovary (CHO) cells are the heavy-duty workhorse in the biopharmaceutical industry [1]. There are many reasons choosing CHO as main commercial protein manufacturer, such as safety considerations, easy transfer of heterogeneous gene into CHO genomes, adaption to serum free media, pretty fast and robust growth of cells and the ability to express recombinant proteins with human-like post-translational modifications [2-4]. Establishing CHO expressing cell line through traditional methods is time-consuming and labor-intensive work mainly because of the unwanted phenotypic heterogeneity caused by position effect [5, 6]. Position effect which refers to the influence of the chromosomal location of a gene on its activity was proposed for over 90 years [7]. It was reported to have profound influence on genomic engineering [8]: previous researches revealed certain genomic domains could exert a general activating or attenuating influence on the embedded genes for protein expression level [9]. Indeed, position effect was the main reason causing the instability issue in traditional cell line construction process [6]. Site-specific integration (SSI) of transgenes into stable hot spots, however, were generally considered to be able to overcome such phenotypic heterogeneity caused by position effect and thus could maintain long-term expression stability [1, 3, 6, 10, 11].

Although targeting gene of interest (GOI) into stable hot spot looks promising, to discover the hot spot could be challenging. The stable hot spot information open to public is rare probably due to commercial value considerations. Normally, if researchers already had highly expressed cell lines, targeted locus amplification (TLA) could be applied to identify new integration sites. TLA method could selectively amplify and sequence entire genes on the basis of the crosslinking of physically proximal sequences; and thus, it could be applied in discovering the transgenes insertion sites [12]. After identifying hot spots, researchers were required to further screen out the final hit by targeting GOI into different spots. If researchers had no expressing cell lines with good performance, the random integration screening methods by fluorescence or productivity of other reporter gene such as β-galactosidase might be applied.

In this study, we used a new method to identify potential targeting sites based on lentivirus infection combined with Zsgreen1 protein screen. One spot specific information was presented in this article and the corresponding cell line model was passaged over extensively to test this insert’s stableness. Finally, the cell line model was adapted to suspension culture and was explored for its potential in industrial application. In summary, the method mentioned here provided researchers a new way to discover their own stable hot spots for SSI related study.

## 2. Materials and Methods

### 2.1. Lentivirus Construction and titer calculation

The reporting lentivirus were constructed as follows: HEK-293T cells were seeded in T75 flask (Corning, Corning, NY) one day before being transfected by 3 plasmids pSPAX2 (10 μg), pMD2G (5 μg) and pHBLV-CMV-Zsgreen1 (10 μg) by using 75 μl Lipofiter (Hanbio, Shanghai, China). Supernatants were collected after 48-72 hours transfection and were centrifuged at 82700 xg for 2 hours. Thus, the lentivirus was finally obtained and was stored in −80 ℃ freezer.

The titer calculation method was as follows: HEK-293T cells were seeded in 96-we plate and cells within each well were infected with serial diluted lentivirus. Three days later, we chose the well that had 10∼30% rate of green fluorescence cells to determine lentiviral titer. The titer calculation function was as follows: Titer (TU/ml) = cell numbers * fluorescence rate * 1000 / original lentivirus volume used (μL).

### 2.2 Cell culture, lentivirus infection and stable cell line construction

CHO-K1 cells were obtained from ATCC and were cultured in Ham’s F12K media (Thermo Fisher Scientific, Waltham, MA) supplemented with 10% FBS (Thermo Fisher Scientific) at 37℃, 5% CO_2_ incubator. Cells were seeded on 6-well plate one day before lentivirus infection. In the next day, lentivirus was thawed on ice and was mixed with 1ml fresh medium. Old medium within well was replaced by this lentivirus mix. Cells were infected by this lentivirus mix for 4 hours. Another 1 ml of medium was added to the plate after 4 hours. In the next day, all medium was replaced by totally new fresh medium. After 72 hour’s infection, pools were single cell sorted and seeded into 96-well plate by FACS based on fluorescence intensity.

### 2.3 Genome Walking

Genomic DNA was isolated and purified by using NucleoSpin Tissue and NucleoSpin Gel and PCR Clean-Up (Clonetech, Mountain View, CA), respectively. The prepared genomic DNA were then separately digested by three different restriction endonucleases: DraI, SspI and HpaI (Clonetech) at 37℃ overnight. The cutting products were all purified by NucleoSpin Gel and PCR Clean-Up and were ligated to genome walker adaptor (Clonetech) at 16℃ overnight to make libraries. All three libraries were amplified by using Advantage 2 PCR Kit (Clonetech). This primary PCR was carried out by using two-step method (5X 94℃ for 25 sec, 72℃ for 3min; 20X 94℃ for 25 sec, 67℃ for 3min; 1X 67℃ for 7min) with primer sets AP1 and LSP1. The primary PCR products were used as template for the secondary PCR. The secondary PCR reaction condition was the same as the primary PCR except for using another primer sets AP2 and LSP2. The secondary PCR products were all sequenced.

### 2.4 Adaption to suspension culture

The original 2C3 model cell line’s culture media were gradually replaced by serum free media M2+M4 (1:1) (Kangju, Suzhou, China). Cells were transferred to a shaking bottle when % FBS was 0 and were cultured for another 3-4 weeks with the speed of 100 rpm. If cells died and the concentration was less than 10^6^ cell/ml, adherent cells should be added into the bottle to maintain a minimum concentration of 10^6^ cell/ml.

### 2.5 Knock-in cell line construction and PCR verification

Cells were transfected with three plasmids: Cas9, sgRNA targeting the integration site and corresponding donor plasmid (molar ration 1:1:1). For each sample, 4×10^5^ cells were transfected with a total of 3 μg of DNA using Lipofectamine 3000 (Thermo Fisher Scientific). Stable cell pools were generated by adding 5 μg/ml puromycin as selection pressure to each well on day 3. After 14 days of selection, cells were detached by TrypLE (Thermo Fisher Scientific) and were resuspended in PBS. The cell pool was sorted out by MoFloXDP FACS machine (Beckman Coulter, Boulevard Brea, CA) and one cell was seeded per well in 200 μl medium in 96-well plate.

Genomic DNA of cells was extracted from pellets and was used as the templates for the following PCR reactions. All PCR reactions were conducted by using Phantom Max Super-Fidelity DNA Polymerase (Vazyme, Nanjing, China). The specific reaction condition for 5’/ 3’ junction PCR was as follows: 95℃ for 3 min; 30X: 95℃ for 15 sec, 66℃ for 15 sec, 72℃ for 2 min; 72℃ for 5 min. Out-out PCR was carried out by using the following condition: 95℃ for 3 min; 30X: 95℃ for 15 sec, 66℃ for 15 sec, 72℃ for 6 min; 72℃ for 5 min. For the nested PCR, which used out-out PCR products as template; the PCR reaction condition was 95℃ for 3 min; 30X: 95℃ for 15 sec, 65℃ for 15 sec, 72℃ for 2 min; 72℃ for 5 min. 5’/3’ junction PCR products and nested PCR products were sequenced.

### 2.6 FACS sorting and flow cytometry analysis

The adherent 2C3 cell line was resuspended in PBS (GE Healthcare Life Sciences, Logan, UT) and cells were single cell sorte and collected by using MoFloXDP FACS machine (Beckman Coulter, Boulevard Brea, CA). Aherent fluorescent Cells were further analyzed by BD FACCcalibur flow cytometry machine (BD, Franklin Lakes, NJ) to verify the heterogenous gene expression stability of cell pools over passages. For the suspension cells, their fluorescence signal was detected and analyzed by MoFloXDP FACS machine again.

## 3. Results

### 3.1 Reporting lentivirus construction and titer detection

Normally, lentivirus expressed reporter gene when it was integrated into the genome. Here we applied Zsgreen1 as our reporter gene for further research. When lentivirus was successfully constructed, its titer value was estimated based on the supplementary Fig 1 where the fluorescence rate was about 20% and the titer was ∼10^8^ TU/ml based on formula mentioned in method part.

### 3.2 Highly expressed model cell line construction and identification of insert’s site

To identify potential hot spots within CHO-K1 cell line genomes, we set up a high-throughput screening method to achieve the goal. The CHO-K1 cells were infected with lentiviral construct harboring the Zsgreen1 gene driven by ubiquitously expressed cytomegalovirus (CMV) promoter at a low multiplicity of infection (MOI = 0.3) to favor single integration. Fluorescence-activated cell sorting (FACS) were applied to isolate single Zsgreen1-positive cells with high fluorescence intensity. The rate of Zsgreen1-positive cells was 4.2% and only cells ranking within top 10% fluorescence intensity were single cell sorted and seeded in 96-well plates for further expansion. The monoclonal cells were monitored under the fluorescence microscope to eliminate non-stable positive cells and cells in slow growth rate. Those qualified colonies were expanded step by step. The genomic DNA from the viral constructs were extracted and were further analyzed by genome walking method to identify all potential viral integration sites within genome [13-15]. The flow diagram of overall high-throughput screening process was depicted in Fig.1.

**Fig 1.**
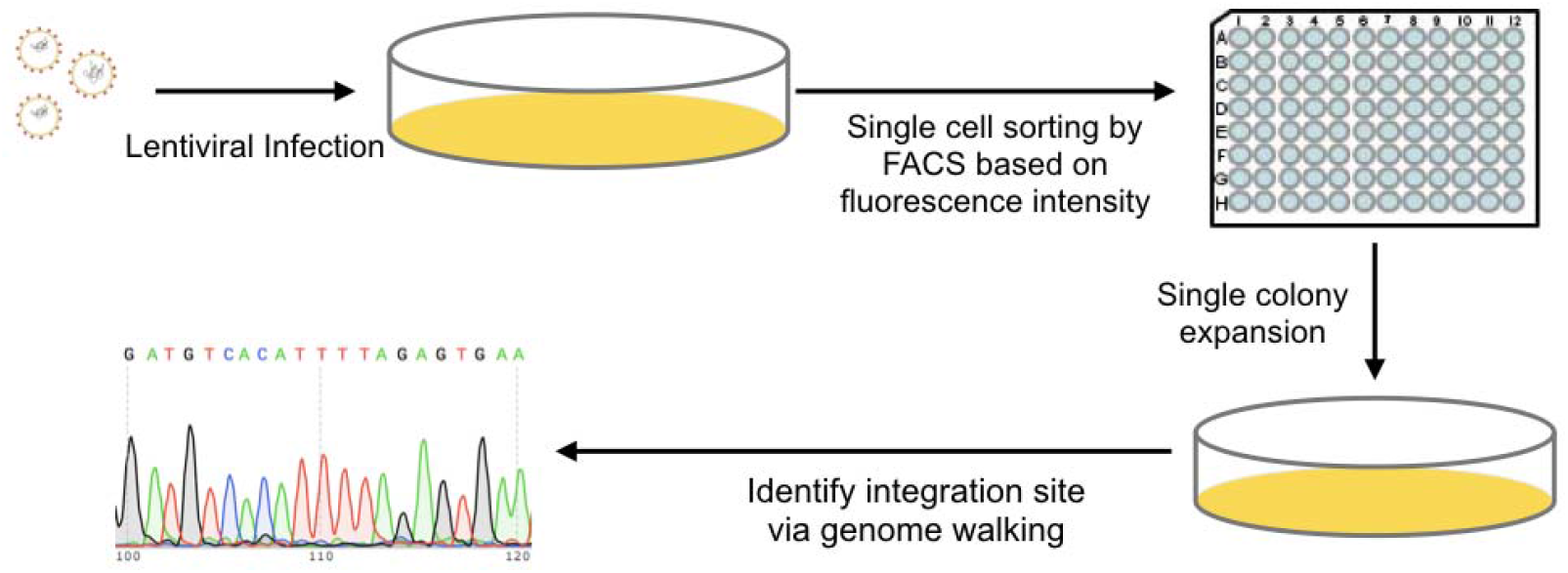
Flow diagram of hot spot discovery by using lentivirus. A lentivirus harboring Zsgreen1 reporter gene was used to infect CHO-K1 cells in low MOI (MOI=0.3). The pool was single cell sorted based on green fluorescence intensity (top 10%). Cells were expanded for over 5 weeks and were undergoing further genome walking analysis.

In this article, only one monoclonal cell line’s (2C3) specific integration site information was presented. The 2C3 cell line’s colony image (Fig.2a) was captured 6 days after FACS sorting: the colony was round shaped, with strong fluorescence signal and containing massive cells which indicated the cells grew well. Thus, this cell line was chosen for later study to identify specific viral integration site. The lentivirus integration site of 2C3 cell lines were identified by using genome walking method.

**Fig 2.**
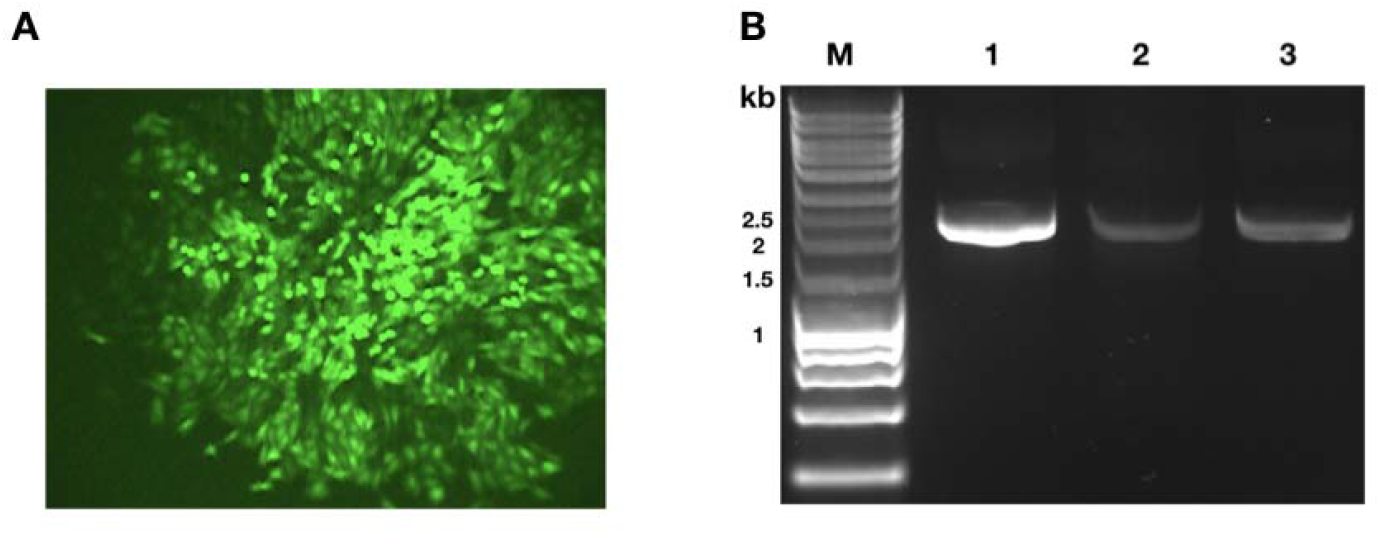
Hot spot identifcation. A. Image of 2C3 cel line captured by fluorescence microscope after 6 days of seeding. B. Lane 1, 2 and 3 corresponded to secondary PCR results of genomes cut by DraI, SspI and HpaI, respectively.

After doing secondary PCR, three samples from different libraries were running on 1% agarose gel (Fig 2b). There was only one band in all three different lanes, indicating only one copy of lentivirus was inserted in 2C3 cell line’s genome. The PCR products were further sequenced by primer AP2. The sequencing results from three different libraries matched with each other (Supplemental Table 1). The hot spot was located at 1235357 within the scaffold NW_006880285.1 analyzed by BLAST at NCBI.

In addition, the hot spot we discovered was located in a copy number variance (CNV) stable region, where CNV value was 2.0 based according to the results of Kaas’s [16]. Hence, this hot spot could be considered as stable from CNV side and was worth of investigation in the further.

By applying this lentivirus-based screening method, we successfully discovered a couple of integration sites which could authenticate its practicability. Moreover, the disclosed 2C3 cell line was required to test its stability in order to evaluate to its potential for industrial application.

### 3.3 Stability assay of adherent model cell line

The stability issue was critical for CHO industry. Based on previous research [17], the unstable expression for CHO cells was related with both genetic factors such as gene copies loss in proliferating CHO cell population and epigenetic factors such as promoter methylation. To further verify the potential of the site for industrial application, we needed to test the model cell lines’ fluorescence signal to make sure it would not disappear over passages. The model cell line was cultured for over 50 passages and fluorescence signal of different cell line passages were detected by flow cytometry. We found the fluorescence rate of cells for both passage 1 and passage 50 were 100% compared to parallel control sample (Fig 3a, 3b, 3c). Hence, the site identified from model cell line could be considered as stable integrating site and it would be worth to further explore its potential for industrial application.

**Fig 3.**
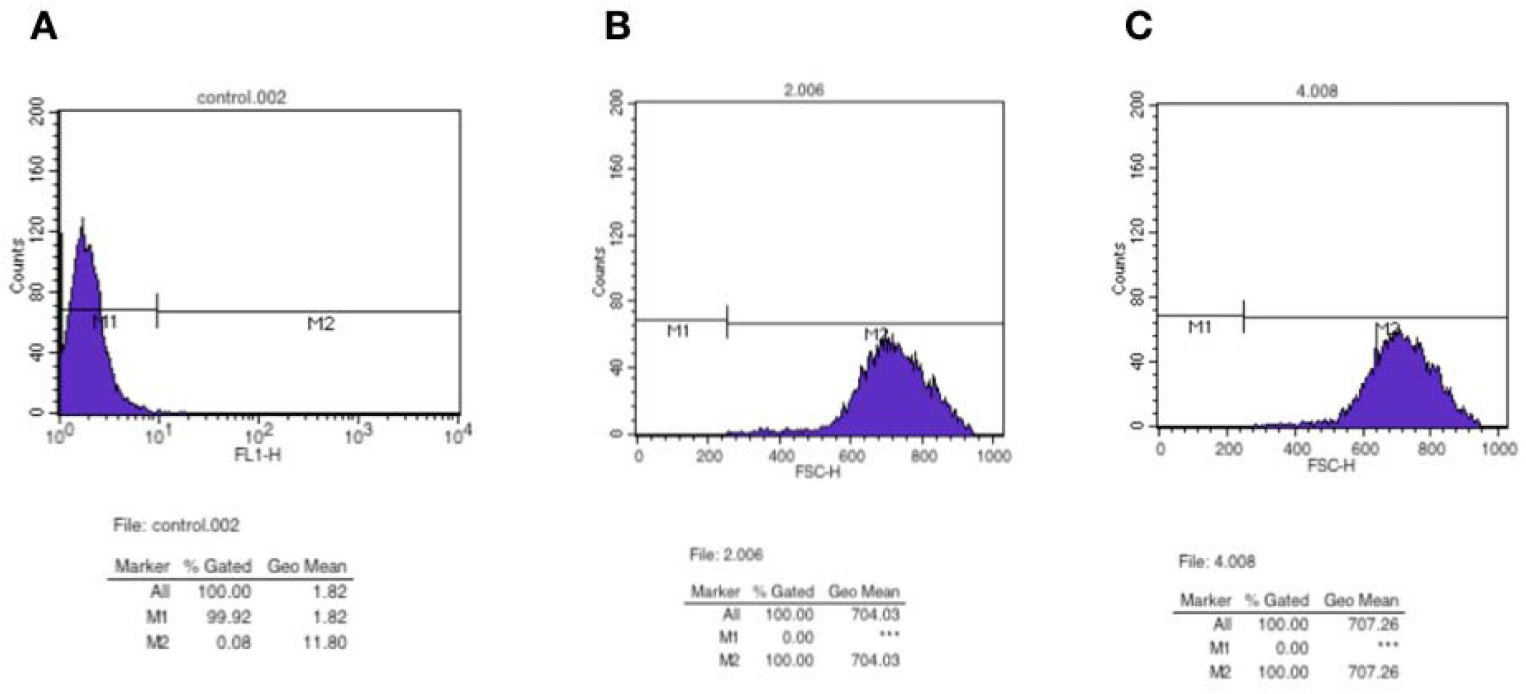
Fluorescence rate of adherent model cell line over passages. A. Fluorescence rate of parallel control sample, 0%. B. Fluorescence rate of 2C3 cell line at passage 1, 100%. C. Fluorescence rate of 2C3 cell line at passage 50, 100%.

The specific parallel control sample was obtained by integrating another non-Zsgreen1 gene into the same spot as 2C3 via CRISPR/Cas9 technology. Here we chose glucagon-like peptide 1 (GLP-1) with human serum albumin fusion protein (NGGH) [18] as the targeting gene. The NGGH donor plasmid was targeted into the spot, and we finally got 3 hits. All three hits were able to be amplified by both 5’ junction and 3’ junction PCR (Supplementary Fig 2b). The molecular weight of all 5’ junction’s amplicons was ∼1.7kb which could match with the design (Supplementary Fig 2a). All 3’ junction’s amplicon’s results were ∼1.5kb and could correspond to the design as well (Supplementary Fig 2a). The 5’/3’ genome-donor boundaries were sequenced to verify precise integration of donor plasmid into the genome. Indeed, the sequencing result verified precise integration of targeting cassette into hot spot locus (Supplementary Fig 2c). Out-out PCR revealed all 3 cell lines were heterozygous and were correctly targeted with the intact target integration unit, generating expected size of amplicons (wild type amplicon: 1.2 kb + target integration unit: 4.7 kb ≈ 5.9 kb; Supplementary Fig. 2d). The out-out PCR products (∼5.9 kb) were purified and were used as template for a series of nested PCR, in order to verify that the correct NGGH sequence was targeted into the genome. Indeed, the sequencing result confirmed complete and correct integration of NGGH gene sequence into the hot spot within genome (Supplementary Table 2).

### 3.4 Adaption to suspension culture and stability assay for suspension cells

CHO cells were normally adapted to serum free suspension culture for potential scale up culture in industry. Hence it was worth to test model cell line’s expression performance after its adaption to serum free suspension culture to further confirm its potential for industrial application. We first adapted cells to suspension. When cell density could be doubled in the next day, the cells could be treated as successful adaption to suspension. Here we found cell density reached 1.98×10^6^ cells/ml in Day 2 while the original cell density was 10^6^ cells/ml by Day 1. Cells were then undergoing dilution back to 10^6^ cells/ml. Again, cell density reached 2.08×10^6^ cells/ml by Day 3. Continuous days of observation verified the adaption of suspension culture successfully (Supplementary Fig 3a). The parallel control sample were also undergoing the same adaption process and the cell density could be doubled every day as well (Supplementary Fig 3b).

Fluorescence rate of 3 different passages of suspension model cell line together with parallel control were all detected by using the FACS machine. We discovered the fluorescence rate of passage 1 was close to 98% after the adaption process when compared to the parallel control sample (Fig 4a). In addition, passage 25 and passage 50 samples both held the fluorescence rate of around 93%∼94% (Fig 4b, 4c). Thus, the process of addition to suspension culture did not significantly impact the fluorescence rate of model cell line. These results further verified the stableness of this hot spot and revealed its potential for industrial application in the future.

**Fig 4.**
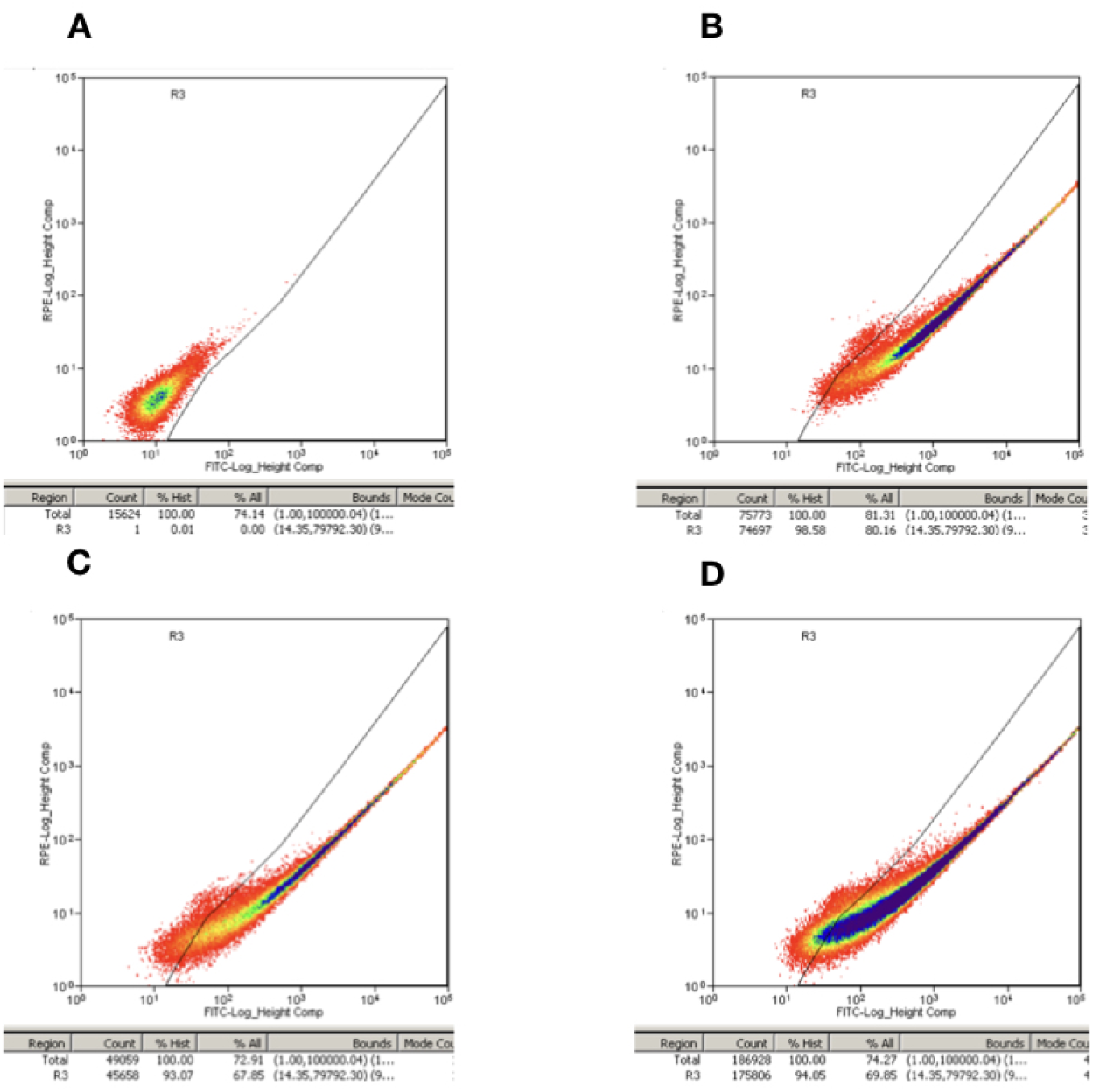
Fluorescence rate of model cell line after being adapted to suspension culture over passages. A. Parallel control sample was used to set up the threshold line to measure fluorescence rate of model cell line. B. Fluorescence rate of model cell line at passage 1. C. Fluorescence rate of model cell line at passage 25. D. Fluorescence rate of model cell line at passage 50.

## 4. Discussion

As CRISPR/Cas9 based knock-in technology became mature, new cell line construction method, SSI of transgenes into the hot spot, became promising in the future. This article provided an easy, high-throughput and low-cost way to identify new hot spot to achieve such new cell line construction method.

To identify new stable hot spots by using lentivirus with fluorescence tag had many advantages. First, integrated form of HIV-1 DNA is traditionally considered to be responsible for viral gene expression [19]. Thus, it provided a good way to link chromatin position with inserts expression level. Plasmid based screening, however, might potentially be interfered by transient expression and thus was not chosen here. Second, the Zsgreen1 reporter gene could achieve high-throughput screening by using FACS compared to other regular reporter gene such as β-galactosidase. Third, the hot spot identification method can be more achievable for researchers compared to TLA method, because this new technology was very complex and expensive. Finally, the fluorescent model cell line itself could be a good tool for researches to do other study, such as media optimization or genetic screening based on CRISPR/Cas9 technology to further improve expression level [20-22].

Stableness was critical for biopharmaceutical industry. Normally, CHO genome was considered as unstable, particularly when cells were under exposure to stress [2]. Thus, traditional cell line construction method by using random integration way cannot guarantee stable cell lines every time [17]. This is because some GOIs were inserted into unstable regions of genome. We proved the hot spot we discovered as stable one from different aspects. The model cell line 2C3 displayed great stableness for over 50 passages (Fig 3). Moreover, when model cell line was adapted to suspension culture, which was great living environment shift for cells, almost all cells kept holding green fluorescence signal (Fig 4b). Additionally, these suspension cells did not lose any fluorescence for another 50 passages (Fig 4c-4d). Interestingly, all these lab data can meet with the stableness predictions based on CNV value [16]. Before being widely applied in biopharmaceutical industry, integration of different GOIs into this stable hot spot to further verify its stableness should be conducted.

Position effect can strongly affect the success of genomic engineering [8] and thus its mechanism would have great commercial value. However, our knowledge in this field was still limited [23, 24]. Current advances in Hi-C technology makes it feasible to investigate the three-dimensional (3D) structure of nuclear organization from different resolutions, including A/B compartments, topological associating domains (TADs) and DNA looping interactions [25] which could help us understand how position effect affects gene expression from different resolutions. Knowledge in this field might potentially improve the production of biopharmaceuticals and enhance our understanding of the mechanism of hot spot’s working way.

In summary, we established one simple and efficient screening method to identify new hot spot within CHO genome and further verified the stableness of one hot spot we discovered. This method will be convenient to researchers to discover their own stable hot spot and apply SSI to construct new expression cell lines in the future.

## Acknowledgement

This work was supported by open lab platform at School of Life Sciences in Fudan University and School of Pharmaceutical Sciences in Jiangnan University. The authors thank Dr. Daru Lu at Fudan University and Dr. Helene F. Kildegaard at Novo Nordisk Foundation Center for Biosustainability, Technical University of Denmark sharing their plasmid. The authors thank Suqin Shen and Longjiang Liu at national key lab platform in Fudan University for the assistance in FACS.

This work was supported by China National Natural Science Foundation Grants 30970029 (to HZ Li.); Ministry of Science and Technology of China Grant 2015AA020802 (to J Jin); National major new drug discovery program of China Grants 2018ZX09738004 (to J Jin).

## Author contributions

ST Zhou, HZ Li, and J Jin designed research; ST Zhou, XF Ding, L Yang, and J Jin, performed research; ST Zhou, Y Chen, HZ Li, and J Jin analyzed data; ST Zhou, XH Gong, and J Jin wrote the paper.

The authors declare no conflict of interest.

